# cSTAR analysis identifies endothelial cell cycle as a key regulator of flow-dependent artery remodeling

**DOI:** 10.1101/2023.10.24.563764

**Authors:** Hanqiang Deng, Oleksii S. Rukhlendo, Divyesh Joshi, Xiaoyue Hu, Philpp Junk, Anna Tuliakova, Boris N. Kholodenko, Martin A. Schwartz

## Abstract

Fluid shear stress (FSS) from blood flow is sensed by vascular endothelial cells (ECs) to determine vessel stability, remodeling and susceptibility to atherosclerosis and other inflammatory diseases but the regulatory networks that govern these behaviors are only partially understood. We used cSTAR, a powerful new computational method, to define EC transcriptomic states under low shear stress (LSS) that triggers vessel inward remodeling, physiological shear stress (PSS) that stabilizes vessels, high shear stress (HSS) that triggers outward remodeling, and oscillatory shear stress (OSS) that confers disease susceptibility, all in comparison to cells under static conditions (STAT). We combined these results with the LINCS database where EC transcriptomic responses to drug treatments to define a preliminary regulatory network in which the cyclin-dependent kinases CDK1/2 play a central role in promoting vessel stability. Experimental analysis showed that PSS induced a strong late G1 cell cycle arrest in which CDK2 was activated. EC deletion of CDK2 in mice resulted in inward artery remodeling and both pulmonary and systemic hypertension. These results validate use of cSTAR to determine EC state and in vivo vessel behavior, reveal unexpected features of EC phenotype under different FSS conditions, and identify CDK2 as a key element within the EC regulatory network that governs artery remodeling.

## Introduction

The vascular system has evolved to provide adequate circulation to the tissues in the face of changing demand due to tissue growth, regression, and changes in metabolic activity. An important aspect of this homeostatic regulation involves sensing of fluid shear stress (FSS) from blood flow by vascular endothelial cells (ECs)^1,2^. ECs encode a FSS set point such that sustained FSS above or below this level triggers vessel outward or inward remodeling to increase or decrease lumen diameter and restore FSS to the initial level^3,4^. Available evidence suggests that FSS near the set point activates signaling and gene expression programs that stabilize the vessel, whereas FSS above or below this range destabilizes the vessel to permit remodeling. Failure of these homeostatic mechanisms results in tissue ischemia associated with excessive inward remodeling in coronary and peripheral artery disease. Vascular malformations associated with excessive outward remodeling and vessel fragility are much less common but must also be considered diseases of compromised vascular homeostasis.

Regions of arteries that bend sharply, branch or bifurcate are subject to FSS of lower magnitude and complex changes in its direction during the cardiac cycle, termed disturbed shear stress (DSS)^5^. Artery segments under DSS are susceptible to metabolic and inflammatory stimuli that induce formation of atherosclerotic plaques, a form of pathological vessel remodeling. Interestingly, most plaques are clinically silent due to compensatory outward remodeling by the unaffected parts of the vessel that preserves lumen diameter, often termed Glagov remodeling^6^. Only in late stage, more severely inflamed vessels, does this mechanism fail, at which point the encroaching plaque decreases lumen diameter to cause tissue ischemia.

These ideas can be conceptualized as a set of interlinked FSS-dependent EC states and transitions between them. Physiological FSS (PSS) at or near the set point confers vessel stability with low permeability, low cell proliferation and turnover, and low inflammation. Low FSS (LSS) that triggers inward remodeling is associated with higher EC turnover, higher EC inflammatory activation with recruitment of leukocytes that assist with remodeling, and smooth muscle contraction. High FSS (HSS) triggers outward remodeling that is associated with cell proliferation, inflammatory activation and leukocyte recruitment, and smooth muscle cell relaxation and proliferation. FSS magnitude and patterns thus governs transitions between distinct EC states that drive vessel remodeling, stability or susceptibility to additional stresses that lead to disease.

We have identified a few pathways that show sharp FSS-dependent regulation and that participate in these processes. For example, the Smad2/3 pathway is specifically activated at low FSS (LSS) and is required for vessel inward remodeling^7^. By contrast, the Smad1/5 pathway is specifically activated by physiological FSS (PSS) and signals vessel stability^3,8^. However, our understanding of these crucially important pathways remains highly incomplete. This limited understanding is a major obstacle to developing safe and effective treatments for vascular diseases including coronary, peripheral and cerebral artery disease, vascular malformations, and vascular dementia. These diseases are due to aberrant vessel remodeling that leads to vascular insufficiency and tissue ischemia, or vessel instability, rupture, and bleeding.

Developing improved therapies requires acquisition of a deep understanding of the regulatory networks that define EC state and transitions between states. Recent efforts applied computational approaches, including machine learning (ML) and computational modeling, to identify different cell states and predict and analyze molecular pathways that govern cell trajectories and cell state transitions^9-11^. A systems biology approach, termed cell State Transition Assessment and Regulation (cSTAR), can use a variety of omics data to identify key regulators of cell state transitions and predict experimental outcomes^12^.

FSS potently determines endothelial cell phenotype, controlling expression of several thousand genes. But a systematic analysis of the distinct transcriptomic responses to each of these flow patterns has not been reported. Thus, to gain insight into these processes, we assessed responses of cultured ECs to LSS, PSS, HSS, OSS, and STAT (no flow) conditions. Use of cSTAR accurately separated the transcriptomic patterns of EC states under these conditions and demonstrated the existence of three orthogonal vector axes that precisely characterize EC state transitions. Combining this information with published EC drug perturbation data (LINCS dataset) allowed us to reconstruct a signaling network that controls EC state transitions and suggested that cell cycle plays a central role in these processes. Experimental studies confirmed these predictions, demonstrate that cell cycle position determines pro- and anti-inflammatory susceptibility, and that CDK2 is a major determinant of the stable, anti-inflammatory state in quiescent ECs under physiological shear stress.

## Results

### cSTAR analysis of EC transcriptomic data under flow

Human umbilical vein ECs (HUVECs) were subjected to OSS (0.5±4 dynes/cm^2^), LSS (3 dynes/cm^2^), PSS (16 dynes/cm^2^), HSS (40 dynes/cm^2^) and STAT for 24h. Total RNA was extracted and prepared for RNA-sequencing. The fold-changes in transcriptomic EC responses to different FSS levels compared to control static condition were considered as points in the transcriptomic dataspace (see Methods).

In the initial cSTAR step^12^, we used machine learning techniques, such as support vector machines (SVM), to exploit the high-dimensional omics space and construct separating surfaces between distinct EC states (technically, the SVM algorithm with a linear kernel from the scikit-learn python library was applied to build a maximum margin hyperplane that distinguishes different cell states). Directions in the transcriptomic space are specified by the State Transition Vectors (STVs), which are unit vectors defined as normal vectors to the hyperplanes that separate cellular phenotypic states. After building the STVs, cSTAR calculates quantitative indicators of cell phenotypic states termed Dynamic Phenotype Descriptors (DPDs). The DPD scores describe the phenotypic cell features associated with each STV. These scores are determined by the Euclidean distances between the separating surface and the cell state that are taken with plus or minus sign depending on the STV directions. If all STVs are orthogonal, then transcriptomics changes along either STV change single, distinct phenotypic features that are associated with different DPDs and determined by generally different molecular processes^13^.

We aim to find the STV directions that describe and determine three different EC functions: (1) fluid shear stress sensing, (2) stability (under PSS), inward (LSS) or outward (HSS) vessel remodeling, and (3) abnormal remodeling (OSS). Therefore, we have used the SVM to separate different transcriptomic responses that correspond to distinct fluid shear stress conditions and cell phenotypic responses. Accordingly, the calculated STVs and DPD scores are denoted by different subscripts. The subscript “FSS” corresponds to the separation of LSS and HSS transcriptomic responses, the “remod” subscript corresponds to the separation of PSS vs LSS and HSS taken together, and the “OSS” subscript corresponds to the separation of OSS vs PSS, LSS and HSS taken together. The resulting STV_FSS_, STV_remod_, and STV_OSS_ are mutually orthogonal, suggesting distinct EC transcriptomic and phenotypic responses to perturbations along these axes. Robustness of the EC state classification and the DPD score variability was rigorously tested using cross validation, as we did previously^12^.

DPD scores calculated along each STV (**Figure 1A-B**) reveal that the remodeling scores, as quantified by DPD_remod_, for EC transcriptomes under both OSS and STAT conditions are very close (**Figure 1A**). These remodeling scores are higher than the scores under PSS, where vessels are stable, but lower than the scores under LSS and HSS that induce strong vessel remodeling. We conclude that endothelial cells under OSS and STAT conditions are in a state that is susceptible to remodeling but less so than under LSS and HSS. The findings for OSS correspond well to behaviors in vivo, where regions of arteries under OSS are stable in the absence of additional stresses, but show high susceptibility to inflammatory and metabolic stresses, which result in preferential inflammatory gene expression and ultimately selective formation of atherosclerotic plaques^14^. ECs under STAT do not exist in vivo but the result fits with general observations that PSS reduces inflammatory gene expression and cell proliferation to below the level seen under static conditions^15^.

**Figure 1.**
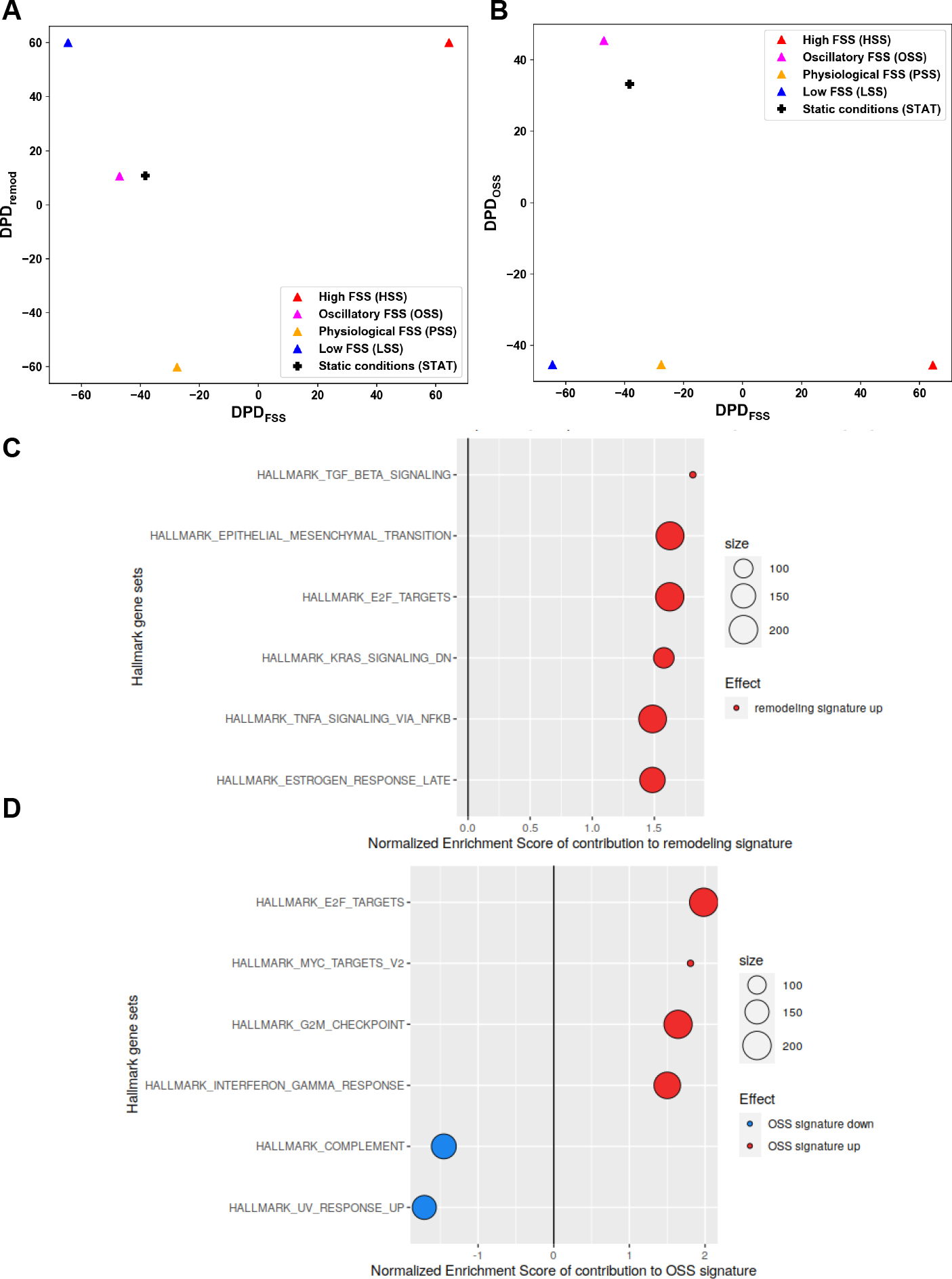
cSTAR analysis of endothelial cell (EC) states under different flow conditions. RNAseq data from ECs under OSS, LSS, PSS, HSS or STAT (static, no flow) conditions for 24h were analyzed and DPD scores calculated for each condition. (A) 2-D plot of phenotypic scores in the DPD_FSS_ and DPD_remod_ plane. (B) 2-D plot of phenotypic scores in the DPD_FSS_ and DPD_OSS_ plane. (C, D) Top GSEA Hallmark gene sets that contribute to the STV_remod_ (C) and STV_OSS_ (D). The components of STVs have been used as the GSEA input for the GSEA hallmark gene set (HM). Positive normalized enrichment score (red) reflects an increase in a molecular process while moving along the corresponding STV, and negative normalized enrichment score (blue) reflects a decrease.

The DPD_FSS_ scores that distinguish LSS from HSS (**Figure 1A-B**) are involved in inward vs. outward remodeling. The DPD_OSS_ scores of OSS and STAT states are also notably close to each other and significantly higher than the scores under all other FSS conditions (**Figure 1B**). Importantly, Figs. 1A and 1B demonstrate that the DPD_FSS_ scores of EC responses under STAT and OSS conditions closely resemble the DPD_FSS_ scores under PSS condition. This suggests that static and oscillatory shear stress conditions predispose vessels to remodeling but without a preference for changing the vessel diameter in a specific direction. Again, this result is consistent with the observation that artery regions under disturbed flow are stable but show high susceptibility to inflammatory and metabolic stimuli^14^.

While both LSS and OSS states have been associated with atherogenic morphological changes^14^, our analysis indicates that these two conditions lead to the emergence of entirely distinct EC states. This distinction is evident in their significantly contrasting positions within the planes representing the phenotypic features of endothelial cell states (see Fig. 1). These disparities arise from substantial differences in the vectors of log-fold changes in gene expression values between LSS and OSS conditions when compared to STAT conditions. To gain a deeper understanding of the differences between the two EC states, we analysed gene contributions to STV_remod_ and STV_OSS_ using GSEA algorithm^16^. As previously described^17^, this approach allows us to discern the transcriptomic features that differentially contribute to the remodeling programs, irrespective of whether the remodeling is directed inward or outward. Our findings revealed two major distinctions (**Figure 1C-D**): (i) The activation of the cell cycle was much more pronounced in the OSS state than in the LSS state. (ii) There were distinct patterns of chemokine expression, with the OSS state associated with the expression of chemokines responsible for macrophage recruitment, while the LSS state was linked to the expression of chemokines responsible for neutrophil recruitment.

### Integrating high-throughput LINCS perturbation data

The derived STVs and DPDs aid in comprehending the differences between endothelial cell states. However, they do not reveal the causal connections within the core networks that regulate these states. To infer these connections, cSTAR relies on perturbation data, which are responses measured in the same omics data space as the STVs and DPDs^12^. Perturbations can be genetic, such as siRNA, or with small molecule inhibitors or activators. The LINCS datasets comprise responses to various drug perturbations but do not encompass different cell states, as all perturbation responses were obtained under a single, namely STAT, condition. Publicly available LINCS datasets are thus insufficient for constructing the STVs and DPDs. However, merging our data with the LINCS perturbation data for HUVEC cells^18^ (GEO accession number GSE70138) offers some insights into how various drug perturbations impact cell states and facilitates the inference of causal connections within the core network governing EC state transitions (**Figure 2A**).

**Figure 2.**
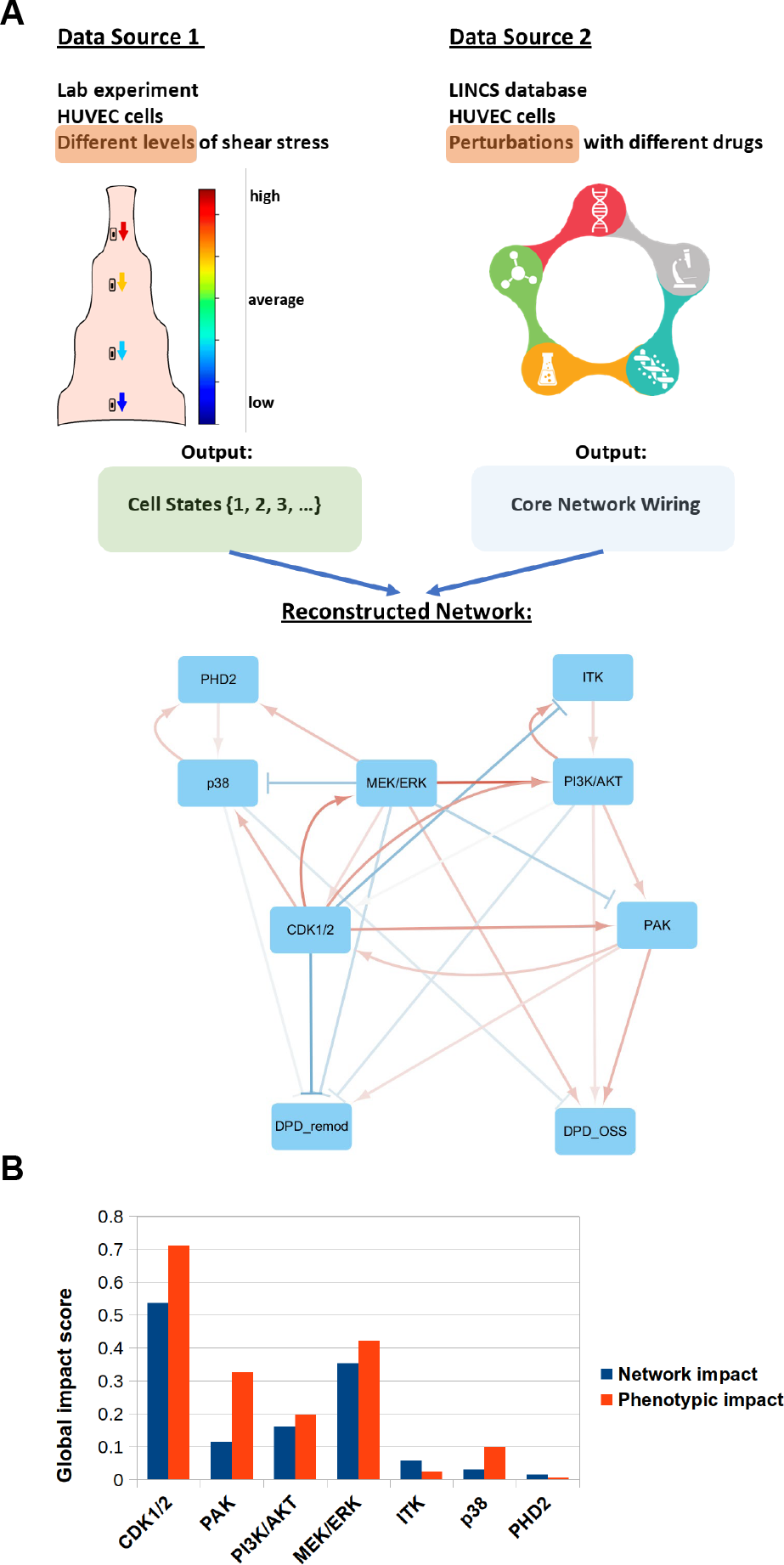
cSTAR reconstruction of the core network driving EC cell fate decisions. (A) Schematic illustrating the data analysis pipeline and the reconstructed core network. RNAseq data from different flow conditions were used to obtain the STVs and perturbation data from LINCS database were used to infer the wiring of the core network and its connection to the phenotypic modules (DPD_remod_ and DPD_OSS_). The resulting inferred network is presented at the bottom. Arrowheads indicate activation, blunt ends indicate inhibition. Line widths indicate the absolute values of interaction strengths. (B) Global impact scores of the effect of perturbation of each core network module on the rest of core network components (blue) and on the cells’ phenotype (red).

The core network components are chosen based on two key criteria. First, these components must exhibit a significant influence on the STVs. Second, their perturbations should yield a substantial impact on the DPDs scores, as outlined in the Methods section. We found that the transcriptomics patterns of the OSS state are close to the patterns of the STAT state (Figure 1). Therefore, we calculated log fold changes of LINCS perturbation data with respect to the DMSO controls and used these data as an input for the cSTAR inference of the core network that controls both OSS and STAT states and EC transitions to other states.

Figure S1 displays the DPD dose responses for drugs primarily influencing DPD_OSS_. It is worth noting that for several targets, such as RAF/MEK/ERK or p38, there exist additional inhibitors that are not depicted in Fig. S1. However, the dose responses to these inhibitors consistently align with those presented in Fig. S1. We find that p38 (TAK-715) and proline dehydrogenase PHD2 (IOX2) inhibitors that upregulate HIF1α increase the DPD_OSS_, whereas RAF/MEK (PD-0325901 and RAF265), MELK/CDK1/2 (OTS-167 and CGP-60474), PAK (PF-03758309), BET bromodomain (JQ1) and ITK (BMS-509744) inhibitors strongly and consistently decrease DPD_OSS_ (Figure S1). According to our framework, drugs that decreasing DPD_OSS_ potentially suppress atherosclerosis related processes, whereas drugs that inhibit p38 and PHD2 (thereby increasing HIF1 expression) potentially promote atherosclerosis. This finding is consistent with reports that HIF1 contributes to inflammatory processes during atherogenesis^19^. Interestingly, the available literature data also suggest that atherogenesis is suppressed by inhibitors of MEK^20^, PAK^21^, and BET bromodomain^22^ proteins. For ITK inhibitors, the situation is less clear. A role for ITK has not been reported but another member of the Tec family, BTK, is a known target in atherosclerosis^23^. Thus, ITK or related kinases are potential anti-atherosclerotic targets. Interestingly, other inhibitors that decrease the DPD_OSS_, differentially affect the DPD_remod_ (Figure S1A). The RAF/MEK and MELK/CDK1/2 inhibitors strongly increase the DPD_remod_, implying that they stabilize the vasculature. By contrast, inhibitors of PAK (except the highest dose), BET bromodomain and ITK/PLCg inhibitors do not significantly change the DPD_remod_ (Figure S1A). These results provide potentially important clues about pathways that regulate EC phenotype, but must be interpreted with caution in the absence of experiments assessing their activation state and function under different flow conditions.

### Inferring causal connections of the core network that controls EC state transitions

Based on their STV contributions and their perturbation impacts on the DPDs, we have selected the components of the core network with the strongest effects. These are: p38, MEK/ERK, PI3K/AKT/mTOR, PHD2, PAK, CDK1/2, and ITK. To infer causal network connections, the changes in the activities of these pathways, kinases and PDH2 that result from drug perturbations need to be determined. The LINCS datasets contain only transcriptomic responses to drug perturbations. The existing methods for identifying enzyme transcriptional signatures and estimating enzyme activities are founded on database knowledge and cover only a few well-studied pathways^12,24^. Therefore, we have developed a data-driven identification of enzyme activity signatures (Methods) that enabled us to not only deduce the drug-induced alterations in the activities of well-known core network modules, such as the MEK/ERK or PI3K/AKT/mTOR pathways, but also for enzymes whose transcriptional signatures are not well-established, such as PHD2, PAK and ITK.

We previously introduced Modular Response Analysis (MRA)^25,26^ and its Bayesian reformulation (BMRA)^27^ as tools to precisely reconstruct causal connections between network nodes, including feedback loop connections, from perturbation data. Each node in the network represents a reaction module, which could be a single protein, gene, or pathway, or any entity defined by its input-output relationships. This enables us to incorporate EC phenotypes, quantified through the DPDs, as network modules. The DPD modules contain information about the performance of the comprehensive cellular network, which is controlled by the core network and plays a critical role in determining endothelial cell phenotypic features.

The network topology is quantified by the connection coefficients, a.k.a, the connection strengths. Each connection coefficient quantifies the immediate impact of a change in a node (A) on another node (B), assuming that the activities of all other nodes are held constant to prevent indirect influence. The connection strengths cannot be directly measured, as perturbations to any node rapidly propagate through the network masking direct, causal connections. We use the BMRA inference, due to its enhanced resilience to noise and its reduced demand for perturbation data, as compared to MRA. Moreover, it offers the flexibility to incorporate pre-existing knowledge about the network in the form of a prior distribution. In BMRA, the inputs consist of the changes in the activities of core network components induced by drug perturbations, along with a prior distribution, which may contain all zeros if no prior knowledge is available. The outputs of BMRA encompass the inferred connections, the associated connection strengths, and their respective confidence intervals^12,27^. Critically, the BMRA algorithm is robust to noise, does not require perturbing all network nodes and outperforms all existing methods of network inference given at least partially correct prior knowledge^27^. Also, BMRA calculates the confidence intervals for each inferred connection, which estimates robustness of the inferred causal connections, including connections to the phenotypic DPD modules^12^.

The inferred causal network connections are presented in Table S1 and Figure 2A. Only two DPD modules, DPD_remod_ and DPD_OSS_, are included in the core network, because drug-induced changes in the DPD_FSS_ scores, which determine the direction, inward or outward, changes in the vessel diameter, cannot be analyzed using LINCS datasets where no SS was applied to ECs.

The causal connections and their strengths quantify the local impact of each node on other nodes, but do not provide insights into the overall, or global changes once a network has equilibrated following a perturbation^25^. These overall changes determine the global node impact. The so-called forward MRA approach allows us to predict the global node impact^28^. This procedure requires the inversion of the inferred network topology matrix, underscoring the significance of each connection within the network for the global changes in the network state. In mathematical terms, the causal coefficient matrix, **r**, tells us how each node immediately affects its neighbors, and the global response matrix, **R**, tells us how each node affects the network regulation globally^28^.

To gain insight into the control of EC remodeling programs, we analyzed both the matrix of the inferred connection coefficients and the matrix of the global node impacts. The causal, local connections show positive feedback loops between the CDK1/2 and ERK, and CDK1/2 and PI3K pathways (**Figure 2A**). Because of these positive feedback loops, these pathways will be activated and deactivated together. As a result, their global phenotypic effects are qualitatively similar (although quantitatively different). These three pathways, CDK1/2, ERK and PI3K, suppress DPD_remod_, suggesting that their inhibition will destabilize vessels to facilitate remodeling. All three pathways also promote DPD_OSS_. Further, PAK directly activates both DPD_remod_ and DPD_OSS_, whereas ITK only weakly affects the DPD nodes. While the pairs of PAK and CDK1/2, and PI3K and ITK are connected by mutual local positive connections, there are also negative connections from ERK to PAK and from CDK1/2 to ITK. This results in differential global responses of PAK and ITK to CDK1/2, ERK and PI3K inhibition. The global impact of PAK activation on the DPD_remod_ is minimal, primarily because of the direct effects of PAK and its indirect effects through CDK1/2 counteract. Thus, global effect of PAK inhibition is suppression of the DPD_OSS_ without affecting DPD_remod_, aligning with reports that PAK inhibition mitigates atherogenesis^21,29-31^.

Besides the interconnected CDK1/2-ERK-PI3K cluster and the subtle regulatory effects of ITK and PAK, there is another cluster of co-activated modules linked by local mutual positive connections: the p38-PHD2 cluster. Whereas PHD2 does not directly connect to the DPD modules, p38 directly suppresses both. The overall effect of p38 activation on DPD_remod_ and DPD_OSS_ is negative, suggesting that p38 suppresses both normal and OSS-activated vessel remodeling. In summary, the primary regulators of signaling nodes, in descending order, are CDK1/2, ERK, and PI3K (**Figure 2B**). The quantitatively assessed global impact on the DPD modules (**Figure 2B**) suggests CDK1/2, PAK, and ERK as the top regulators.

These combined analyses predict that the CDK1/2 module is the most crucial in regulating various SS-induced endothelial cell states (**Figure 2B**). This underscores the pivotal role of endothelial cell cycle regulation in vessel homeostasis. Particularly, our analysis predicts that CDK1/2 inhibition will strongly increase the DPD_remod_ score, thus promoting vessel remodeling. While the roles these regulators play in shear stress signaling must be inferred with caution, they are consistent with numerous studies linking shear stress to cell cycle regulation such that physiological shear stress inhibits while oscillatory or other disturbed shear stress conditions allow greater proliferation^32-38^. We therefore assessed the role of cell cycle regulation in shear stress dependent artery remodeling using experimental approaches.

### Effects of FSS on cell cycle

We first assessed cell cycle state under the different FSS conditions using the FUCCI reporter system that distinguishes cells in early G1, late G1 and S-G2-M. FUCCI is based on expression of the fusion proteins mCherry-hCdt1 and mVenus-hGem that are differentially ubiquitinated and degraded at different points in the cell cycle^39^. During S, G2 and M phases (S-G2-M), only mVenus-Geminin is expressed, resulting in cytoplasmic green fluorescence^40,41^. Following mitosis, geminin is degraded, and cells are non-fluorescent (Go/early G1). As cells progress to late G1, mCherry-Cdt1 is expressed and red fluorescence accumulates. We generated stable FUCCI-expressing HUVECs, subjected these cells to different FSS conditions for 24h and scored cells as unlabeled, red or green. We found that OSS increases cell cycle progression as indicated by an increase in green cells (S-G2-M phases), LSS induces significant early G1 arrest (FUCCI negative), PSS induces strong late G1 arrest (FUCCI red), while HSS increases cells in S-G2-M, indicating greater proliferation (**Figure 3A-B**). To confirm these findings, cells were labelled with the thymidine analog EdU during the last 2h of the 24h flow regimen and stained for this marker to identify cells that synthesized DNA in this interval. EdU labeling demonstrated that OSS induced the highest DNA synthesis, LSS suppressed, PSS strongly suppressed and HSS had little effect on DNA synthesis compared to static (**Figure 3C-D**). These results confirm the findings from the FUCCI reporter and identify an unexpected difference in the nature of cell cycle arrest that depends on flow magnitude. They are also consistent with published findings that cell proliferation is increased under OSS and reduced by PSS^32,35-38^.

**Figure 3.**
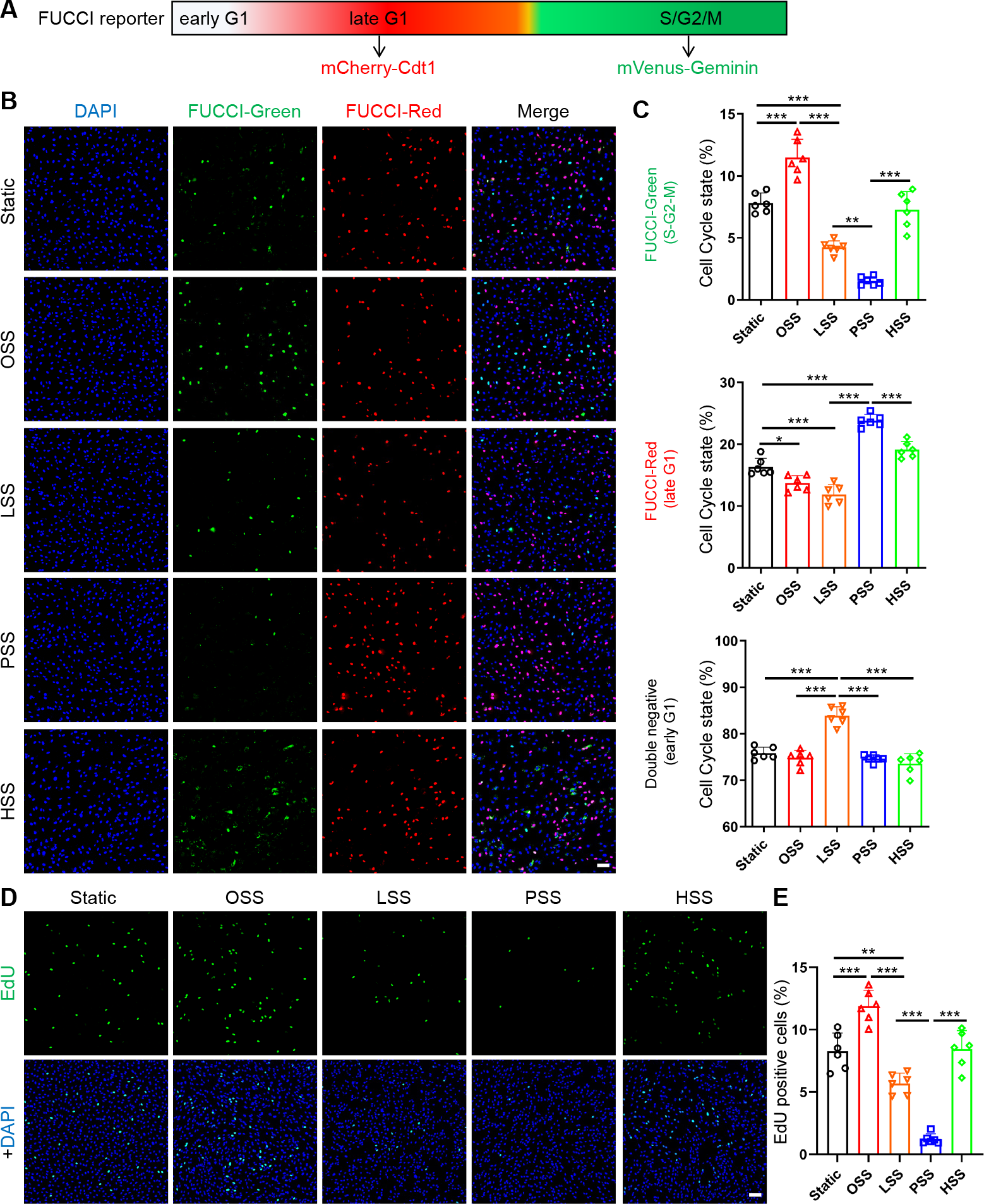
Characterization of cell cycle state under different FSS. (A) Diagram of the FUCCI reporter. (B-C) FUCCI HUVECs were subjected to OSS, LSS, PSS, HSS or STAT for 24 hours. Cells were fixed, stained with DAPI and imaged as described in Methods. Representative images for these conditions shown in (B). Scale bar: 100μm. FUCCI-red, FUCCI-green positive cells and double negative cells were quantified (C). (D-E) EdU assay. HUVECs were subjected to flow patterns as in A, labelled, and stained for EdU as described in Methods. Percent of EdU positive cells was quantified. n=6 experiments. Scale bar: 100μm. \**P* < 0.05, \*\**P* < 0.01, \*\*\**P* < 0.001, ns: not significant, calculated by one-way ANOVA with Tukey’s multiple comparison tests.

### CDK2 activity and function in ECs

Active CDK2 complexed with cyclin E activation is thought to be the major determinant of the late G1-to-S transition^42^. Our previous work showed that the Smad2/3 pathway, an important mediator of endothelial inflammatory gene expression during atherosclerosis and low flow inward remodeling in vascular ECs, is directly inhibited by CDK2^7^. We therefore wished to assay CDK2 activity under these FSS patterns, mainly to determine whether the late G1 arrested state under PSS is associated with active CDK2. For this purpose, we used a reporter construct, DHB-mVenus^43^, whose phosphorylation by CDK2 triggers its translocation out of the nucleus (**Figure 4A-B**). HUVECs were co-transfected with the DHB-mVenus and histone 2B (H2B)-mCherry as a nucleus marker; cells were then subjected to LSS and PSS. We found that PSS increased the cytoplasm/nucleus ratio by more than 10X, while LSS had no significant effect relative to the static, no flow condition (**Figure 4C-D**). CDK2 is thus strongly activated in the PSS, late G1 arrested state but not in the LSS early G1 arrested state.

**Figure 4.**
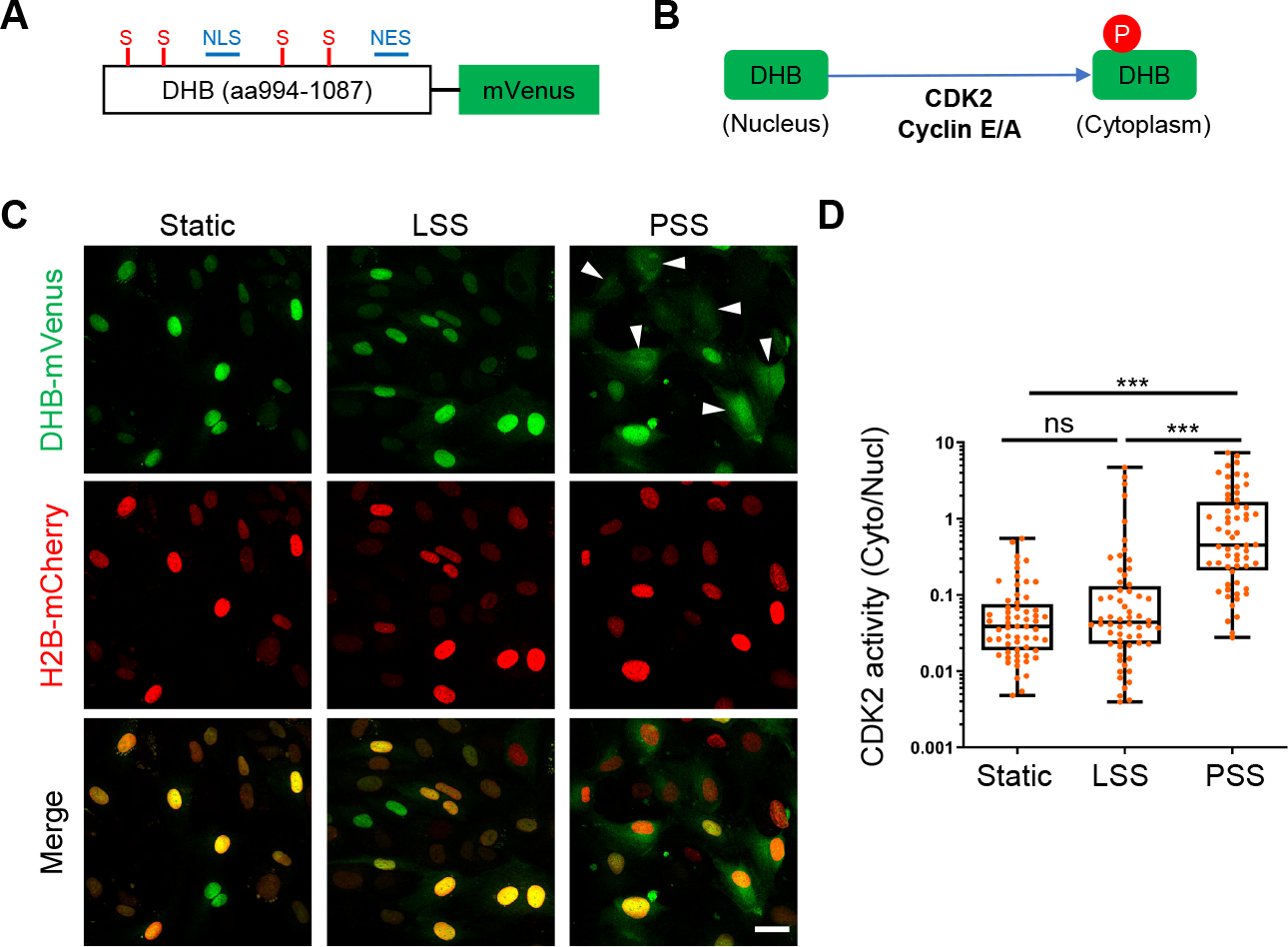
CDK2 activity under FSS. (A) Schematic of sensor. NLS, nuclear localization signal; NES, nuclear export signal; S, serines that are CDK consensus phosphorylation sites. (B) Schematic of CDK2 phosphorylation-mediated translocation of DHB-mVenus. (C-D) HUVECs expressing the DHB sensor were subjected to indicated FSS conditions for 6h. Cells were fixed and imaged as described in Methods, representative images shown in (C). Arrowheads are CDK2 high activity cells. Cell nuclei were identified using fluorescent H2B images to obtain a mask. Ratio of cytoplasmic/nuclear signal was quantified (D), n=60 cells for each group from 3 independent experiments. \*\**P* < 0.01, \*\*\**P* < 0.001, ns: not significant, calculated by one-way ANOVA with Tukey’s multiple comparison.

To further evaluate how CDK2 affects cell cycle state, we performed CDK2 knockdown in FUCCI HUVECs (**Figure 5A**). CDK2 depletion strongly reduced the fraction of cells in S-G2-M and increased early G1 arrest (**Figure 5B-C**).

**Figure 5.**
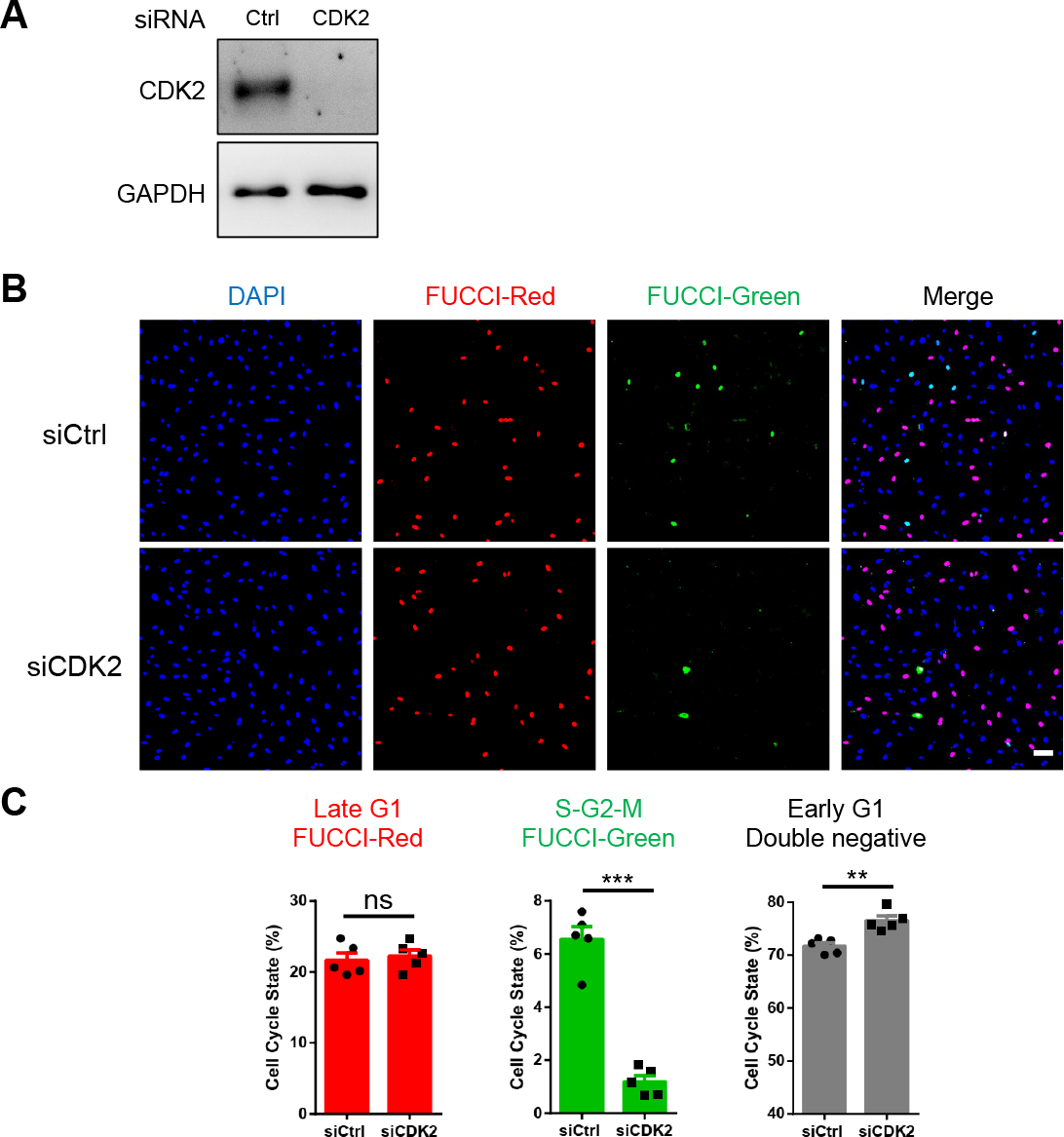
CDK2 depletion induces early G1 arrest. FUCCI HUVECs were transfected with control (siCtrl) or CDK2 (siCDK2) siRNA for 4 days. CDK2 knockdown efficiency was confirmed by Western blotting (A). Cells were fixed and mounted with DAPI. Representative images are showed in (B). Scale bar: 100μm. (C). Quantification of cell cycle state, n=5 experiments. \**P* < 0.05, \*\**P* < 0.01, \*\*\**P* < 0.001, ns: not significant, calculated by two-tailed unpaired t tests.

### Deletion of CDK2 in ECs in vivo

We previously found that a pharmacological inhibitor of CDK2 and related CDKs induced artery inward remodeling and pulmonary arterial hypertension^7^, though this protocol was limited by general toxicity. To investigate the role of CDK2 in vivo using a more specific approach, we crossed CDK2 floxed mice with Cdh5-CreERT2 mice to induce endothelial-specific CDK2 knockout (CDK2 iECKO; **Figure 6A**). Mice were injected with tamoxifen to induce CDK2 iECKO at 8 weeks of age, and Cdk2 deletion confirmed (**Figure 6B**). Mice at 2 months after CDK2 iECKO showed markedly higher right ventricular systolic pressure (RVSP) and less dramatic but significant left ventricular systolic pressure (LVSP) (**Figure 6C-E**). CDK2 iECKO thus induces pulmonary arterial hypertension and to a lesser extent systemic hypertension. To investigate mechanisms, we focused on the pulmonary vasculature where effects were stronger. Immunostaining of smooth muscle actin (SMA) in whole lung sections showed an increase in muscularized small vessels, typical of pulmonary hypertension and indicating artery inward remodeling (**Figure 6F-G**). Masson’s trichrome and H&E staining of lung sections confirmed that arteries in CDK2 iECKO lung became narrowed and even occlusive (**Figure 6H**, Figure S2). Immunostaining of phospho-Smad2 and 3 in lung sections demonstrated their hyperactivation after CDK2 iECKO (Figure S3). CDK2 iECKO thus leads to artery inward remodeling and consequent hypertension.

**Figure 6.**
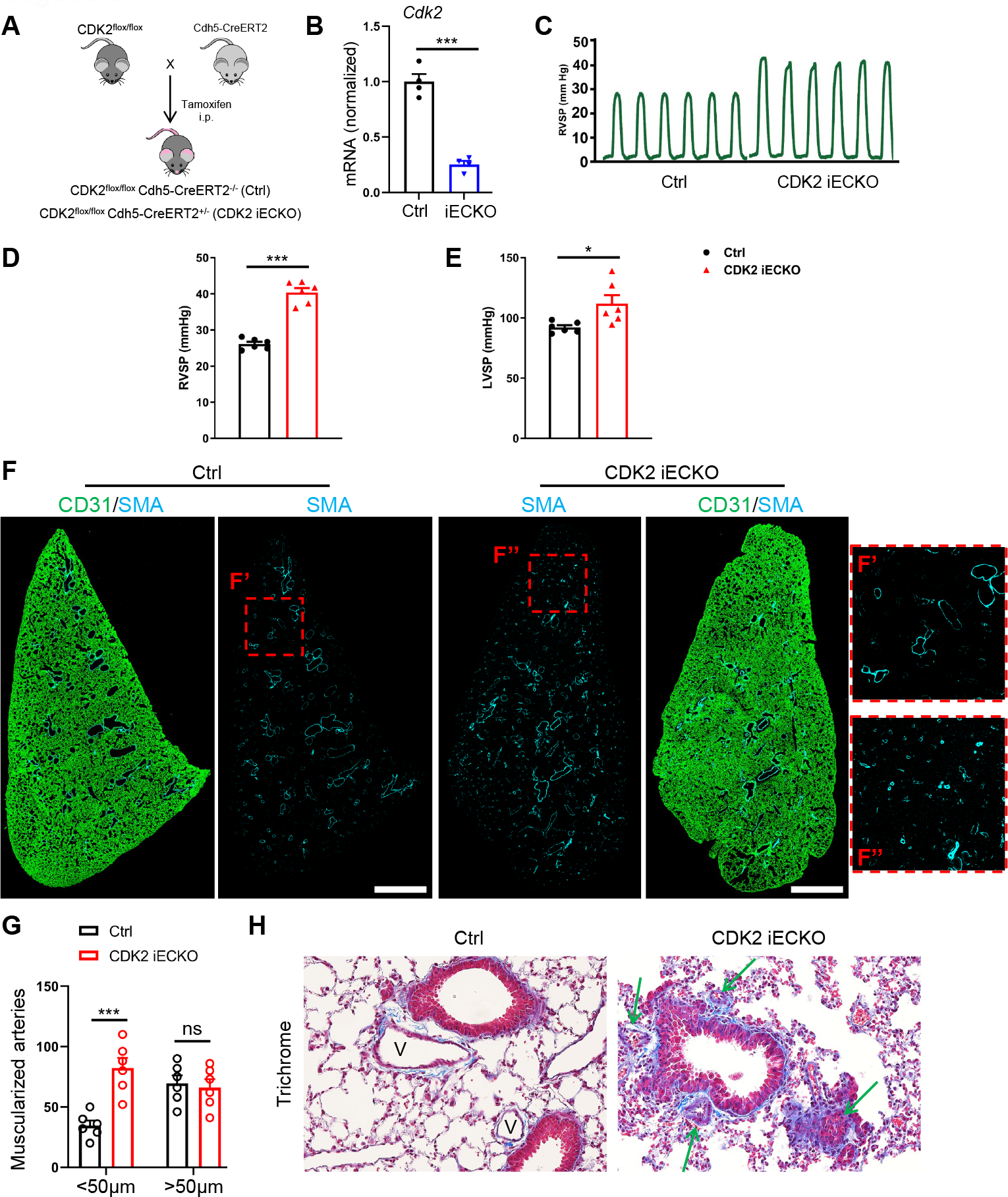
Deletion of endothelial CDK2. (A) Cdh5-CreERT2; CDK2^flox/flox^ (CDK2 IECKO) and CDK2^flox/flox^ (ctrl) mice at 6 weeks of age were injected with tamoxifen. (B) Lung ECs were isolated to check Cdk2 deletion efficiency by Q-PCR. (C-D) Representative and quantification of right ventricle systolic pressure (RVSP) from control and CDK2 iECKO mice. (E) Quantification of left ventricle systolic pressure (LVSP) from control and CDK2 iECKO mice, n=6 mice per group. (F) Representative SMA immunostaining in Ctrl and CDK2 iECKO entire lung sections. Scale bar: 1mm. (G) Quantification of SMA^+^ muscularized arteries (diameter <50μm and >50μm) in Ctrl and CDK2 iECKO lungs. (H) Representative Masson’s trichrome staining of Ctrl and CDK2 iECKO lung sections. V, vessels. Arrows indicate narrowed and occlusive vessels. \**P* < 0.05, \*\**P* < 0.01, \*\*\**P* < 0.001, ns: not significant, calculated by two-tailed unpaired t tests (B, D, E) and two-way ANOVA with Tukey’s multiple comparison (G).

## Discussion

The results reported here lead to three main conclusions of general interest. First, cSTAR analysis of transcriptomic data for endothelial cells under low, physiological, high and oscillatory FSS conditions identifies three distinct, orthogonal axes that describe endothelial states (Fig 1). In keeping with published studies which show that PSS confers vessel stability, ECs under PSS have a low value on the remodeling axis (DMD_remod_) and an intermediate value on the FSS axis (DMD_FSS_), indicating no propensity to remodel in either direction. Interestingly, ECs under both STAT and OSS show intermediate values on the remodeling axis, indicating a less stable state than PSS but well below the actively remodeling LSS and HSS conditions. OSS and STAT conditions have intermediate values on the DMD_FSS_ axis, similar to PSS, again indicating no propensity for inward or outward vessel remodeling. OSS and STAT are also in close proximity on the OSS axis (DMD_OSS_). These findings correspond well to behaviors in vivo, where artery walls under disturbed, atherogenic flow are stable over decades in the absence of additional perturbations but are specifically susceptible to inflammatory and metabolic stresses that lead to formation of atherosclerotic plaques. Consistent with reduced stability, these regions also show amplified rapid responses to inflammatory stimuli, so-called inflammatory priming, whereas regions of arteries under PSS are resistant. The cSTAR analysis of transcriptional signatures thus offers a novel perspective on FSS-driven EC states.

Second, cSTAR analysis of our transcriptomic data integrated with the LINCS datasets, which describe responses of ECs without flow (STAT) to a library of compounds, identified potential causal connections between multiple pathways that determine vessel stability, remodeling and disease susceptibility. This analysis identified a causal network linking CDK1/2, Erk MAP kinases, PI3K/Akt, PAK, PHD2/HIF1α and ITK that is likely to play a major role in determining EC state. In this model, CDKs, Erk and PI3K suppress remodeling, i.e., confer stability whereas PHD2 and ITK appear to promote remodeling. These outcomes fit well with the known ability of Erk1/2 to promote arterial phenotype which is linked to stability^44^ and PI3K/Akt to promote EC survival and activate eNOS, a major anti-inflammatory, anti-thrombotic pathway^45^. It also fits well with the known pro-inflammatory role for HIF1α and PAK^19,21,29-31^. ITK is not well studied in endothelial cells, and it is unclear whether ITK itself or another TEC family kinase was the drug target in the LINCS studies^23^, thus conclusions about the role of ITK or homologs await further work.

Third, based on the finding that CDK1/2 appeared to play a central role in suppressing DMD_remod_, analysis of EC cell cycle state using the FUCCI reporter system revealed that LSS suppressed cell proliferation via early G1 arrest, while PSS suppressed more strongly but via late G1 arrest. We also found that CDK2 is active in the late G1 arrested state and that its knockdown further decreased cell cycle progression but shifted cells into early G1. In vivo EC deletion of CDK2 resulted in pulmonary and systemic hypertension with inward remodeling of small arteries, providing strong evidence that CDK2 is a major component of the network that stabilized arteries under PSS. Indeed, its deletion is sufficient to switch ECs to an early G1 arrest and initiate inward artery remodeling. This effect is evidently not due to cell cycle arrest per se but rather indicates that early vs late G1 arrest induces highly distinct states. The cSTAR analysis of LINCS data (Fig 2) suggest that CDK2 influences the activity of other mediators (Erk, PI3K), creating positive feedback loops. Thus, CDK2 may function both directly, for example, via phosphorylation and inactivation of Smad2/3, and indirectly through additional mediators such as Erk and PI3K.

Taken together, these results strongly validate the cSTAR approach as a means to elucidate the role of FSS conditions on EC state as a driver of vessel remodeling. They also lead to three important and novel conclusions concerning the nature of FSS-regulated EC states, elucidation of a potential regulatory network (Fig 2), and the role of CDK2 in determining vessel remodeling. However, these findings represent only the first step. Complete elucidation and validation of the FSS-dependent regulatory network will require the type of perturbation analysis described in the LINCS database with cells under the different FSS conditions. Doing so offers the opportunity to fully define the pathways and genes that control artery remodeling and identify therapeutic targets to treat vascular disease.

## Materials and Methods

### Animals

*CDK2*^*fl*/*fl*^ mice and *Cdh5-CreER*^*T2*^ mice were previously described^46,47^. All mice in this study were on the C57BL/6 background. To induce gene deletion in adults, mice were injected intraperitoneally with 1.5mg tamoxifen (Sigma, T5648) for five consecutive days. All mouse protocols and experimental procedures were approved by Yale University Institutional Animal Care and Use Committee (IACUC).

### RV and LV hemodynamic measurements

RVSP was measured with a 1.4F pressure transducer catheter (Millar Instruments) and LabChart software (ADInstruments) as described^48^. Briefly, mice were anesthetized with 2% isoflurane and the catheter was inserted through the right jugular vein into the right ventricle. For LVSP, the catheter was inserted through the carotid artery into the left ventricle. RVSP and LVSP were all recorded and analyzed with LabChart software (ADInstruments).

### Cell culture and siRNA transfection

HUVECs (Human Umbilical Vein Endothelial Cells) were obtained from Yale Vascular Biology and Therapeutics tissue-culture core laboratory at Passage 1. Cells were maintained in EGM2 Endothelial Cells Growth Media (Lonza, CC-3162) and used for experiments between P2 and P5. SiRNA transfection was performed with Opti-MEM medium (ThermFisher) and Lipofectamine RNAiMAX (Invitrogen). ON-TARGET plus Smartpool siRNAs from Dharmacon were used against human CDK2 (L-003236-00-0005).

### Generation of FUCCI HUVECs

HEK293T cells were transfected with pBOB-EF1-FastFUCCI-Puro (Addgene plasmid #86849) and packaging plasmids using lipofectamine 2000 (Thermo Fisher Scientific, 11668019) according to the manufacturer’s instructions. Supernatants containing lentivirus were collected 48 h after transfection and passed through a 0.22μm filter. Primary HUVECs were infected with lentivirus for 24h, then replaced with EMG2 Endothelial Cells Growth Media. Cells were selected and passaged in puromycin (1μg/ml, Sigma P9620).

### Shear stress

HUVECs were seeded on tissue culture plastic slides coated with 20μg/mL fibronectin for two hours at 37°C and grown to confluence. Shear stress with a calculated intensity of 1 dyn/cm^2^ to 40 dyn/cm^2^ was applied in parallel flow chambers as described^7,49^.

### EdU cell proliferation assay

HUVECs were seeded on tissue culture slides, grown to confluence in EGM2 medium, and subjected to shear stress as indicated. After 22h, a 2X working solution of EdU in complete EGM2 medium was added to cells, final concentration 10 μM. 2h later, cells were collected and fixed. EdU DNA synthesis was detected following manufacturer’s Click-iT EdU imaging kits (Invitrogen, C10337).

### Immunohistochemistry

Samples were fixed in 4% paraformaldehyde (Electron Microscopy Sciences) overnight at 4°C and incubated with 30% sucrose (Sigma) solution in PBS overnight at 4°C. Then samples were embedded in OCT medium (SAKURA) and 8-10μm sections were cut in a cryostat (Leica). For immunohistochemistry, samples were incubated in blocking buffer (5% donkey serum, 0.2% BSA, 0.3% Triton X-100 in PBS), followed by incubation with primary and secondary antibodies diluted in a blocking buffer. Images were taken using the SP8 confocal microscope (Leica).

### RNA isolation and quantitative real-time PCR

RNA was extracted from cells with RNeasy Plus Mini Kit (QIAGEN) according to the manufacturer’s instructions, and reverse transcription performed with the iScript Reverse Transcription Supermix for RT-qPCR (BIO-RAD). Then cDNA was amplified by real-time PCR with iQ SYBR Green Supermix (BIO-RAD). The expression of target genes was normalized to expression of housekeeping gene *GAPDH*. Primer sequences were listed as below: mGapdh (5’-AGGTCGGTGTGAACGGATTTG-3’, 5’-TGTAGACCATGTAGTTGAGGTCA-3’); mCdk2 (5’-CCTGCTTATCAATGCAGAGGG-3’, 5’-GTGCTGGGTACACACTAGGTG-3’).

### Computational methods and analysis

Total RNA was extracted from HUVECs subjected with OSS, LSS, PSS, HSS, and static for 24h. Total RNA was quantitated by NanoDrop, and RNA integrity number value was measured with an Agilent Bioanalyzer. Samples were subjected to RNA-sequencing using Illumina NextSeq 500 sequencer (75bp paired end reads). The base calling data from sequencer were transferred into FASTQ files using bcl2fastq2 conversion software (version 2.20, Illumina). To analyze, EC responses to fluid shear stress (FSS), the control condition is selected as no treatment, static condition (STAT). However, NT siRNA have been added to some conditions, but not to every condition, in particular H-FSS was analyzed without the NT siRNA treatment. We performed batch correction manually. First, using DeSeq2 (R package for the analysis of RNAseq data) we determined the differentially expressed genes (DEGs) comparing the conditions L-FSS, P-FSS, O-FSS and knockdown series with the corresponding STAT conditions, where all conditions were treated with NT siRNA. To account for the absence of NT siRNA treatment under H-FSS and the corresponding STAT condition, we added the vector of log-fold changes between no treatment, STAT and NT siRNA, STAT, which was constructed using comparison series, to the high FSS. Therefore, all conditions analyzed below were either treated with NT siRNA or corrected by adding the vector of log-fold changes between no treatment, STAT and NT siRNA, STAT.

### STVs and cell states

The LINCS database contains a large number of transcriptomic responses to different drug perturbations, however, such datasets do not contain cell states. In the cSTAR approach, the STV determines the direction in the data space that corresponds to a cell state transition. Thus, the STVs must be determined from different data. We have built the STVs, which are defined as normal vectors, ***n***, to each separating hyperplane, using our data on HUVEC transcriptomic responses to different FSS magnitudes.

To make the STV transferrable between different datasets, the STVs must be built in the space of log fold changes (LFCs). The reference state, with respect to which LFCs are calculated, is STAT or HUVEC cells in the LINCS database. To estimate phenotypic changes brought about by drug perturbations, we calculated the dot products of the vectors STV_remod_ and STV_OSS_ and the vectors of log fold changes obtained from L1000 data and plotted dose responses. Using LINCS perturbation data, we can calculate the change in the corresponding DPD for each perturbation. Based on these assumptions, we calculate dot products of LFCs from LINCS (***x***_***j***_) with the STV (***n*)**, to find element of the global response matrix (***R*** _*DPD*,*j*_), for the j-th perturbation corresponding to the DPD module, as follows

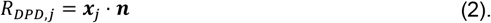

It should be noted that expression (2) implies that for control data point (DMSO, no treatment, NT siRNA, etc.) the *R*_*DPD*_, *j* is equal to zero (or close to zero), because te perturbation vector ***x***_*j*_ is close to zero.

### Global responses of core network modules

If a direct measurement of the core network module activity can be done, finding *R*_*ij*_ elements for the core network modules is straightforward. This is the case for genetic perturbations, such as siRNA, TALE, etc, and the following up measurements of transcriptomics responses. Then, the LFC in the expression of the gene i in response to the j-th perturbation can be used for *R*_*ij*_ value.

In case when the direct measurement of the i-th node activity is not available, we suggest the following procedure. The activity of the of the i-th module after j-th perturbation (*f*_*ij*_) is calculated as linear function of transcriptomic LFCs (*x*_*kj*_) as

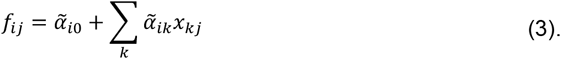

Coefficients 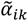, which are used to infer activity of i-th module, can be found using linear regression over function *g* which describes response of enzyme activity to an inhibitor. In the simplest case of monomeric target this function is expressed as follows

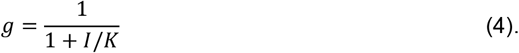

Here *I* is the inhibitor concentration and *K* is IC50 dose for this inhibitor, which must be found in literature. If multiple inhibitors of the same target were applied, the function coefficients 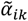 must be found by regressing function *g* over all doses of all inhibitors applied. The coefficient 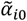 is logically to set to 1 for all modules, because for the control (e.g. DMSO control) data points, all LFC, *x*_*kj*_, are close to zero, while value of the function ***n***is equal to 1.

To satisfy the MRA modular insulation condition [doi: 10.1038/s41540-019-0096-1], the vectors of coefficients 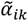 (excluding the intercept coefficient 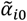) for different modules should be mutually orthogonal. Additionally, we have penalized the negative pathways activities and added Lasso regularization to minimize the number of potential solutions and ideally find the unique solution. Thus, we infer the coefficients 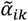 by finding the global minimum of the following function.

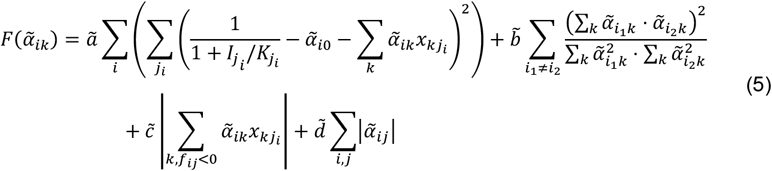

Here the first term describes the linear regression over enzyme activity for each module, the second term introduces the orthogonality condition, the third term penalizes the negative pathway activities and the forth term introduces Lasso regularization. The hyper-parameters, 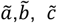 and 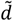, provide a means to balance the priority among different terms when it is not possible to satisfy all conditions simultaneously.

Taking into account that 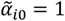, ∀*i*, expression 5 reads

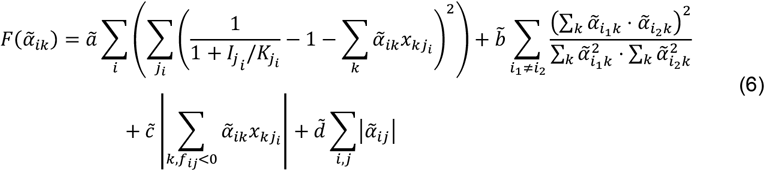

When coefficients 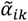 are determined, the elements of global response matrix *R*_*ij*_ for core network modules can be calculated from Eq. 3 as follows,

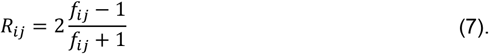

Here we imply that for control data point *f*_*ij*_ *=1*.

It is important to note that while the STV, denoted as ***n***, is normalized to have a unit length, the vectors 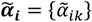 do not have unit lengths and must not be normalized; otherwise, they will not meet the regression conditions.

Completing the entries, *R*_*ij*_, for both the core network modules and the DPD module(s) marks the completion of the transcriptomic data preparation for the BMRA step in cSTAR.

All codes for computational analysis will become publicly available upon journal publication.

### Western Blotting

Cells were harvested and lysed for 30 min on ice in RIPA buffer (Roche) containing complete mini protease inhibitors (Roche) and phosphatase inhibitors (Roche). Centrifuge at 14000 rpm for 10 min at 4°C, transfer cell supernatant to new 1.5 ml EP tubes, and boiled with 4X loading buffer (250mM Tris-HCl pH 6.8, 8% SDS, 40% glycerol, 20% β-mercaptoethanol, 0.008% bromophenol blue) at 99°C for 5 min. Cell lysates were resolved by a 4-15% CriterionTM TGXTM Precast Gels (BIO-RAD) SDS–PAGE electrophoresis, and blotted onto a polyvinylidene difluoride (PVDF) membrane (Millipore). The blotted membranes were blocked with 5% non-fat milk and then incubated with specific antibodies diluted in 5% BSA using a standard immunoblotting procedure and detection by ECL (Millipore). ImageJ was used for densitometry quantification of western blot bands.

### Mouse lung ECs isolation

Mouse lungs were collected and digested in a solution of 2mg/ml collagenase (Sigma). Cell suspension was then filtered through 70μm sterile cell strainer (Falcon). ECs were isolated using magnetic beads anti-rat IgG (Invitrogen) coated with rat anti-mouse CD31 antibody (BD Biosciences). Cells were lysed for RNA extraction with PicoPure RNA isolation kit (Applied Biosystems) according to the manufacturer’s instructions.

### Antibodies

We used the following antibodies for immunohistochemistry (IHC) and immunoblotting (IB): Rat anti-Mouse CD31 (BD 550274; IHC 1:200), GAPDH (Cell Signaling 5174S; IB 1:2000), p-Smad2 Ser465/467 (Millipore AB3849-I; IHC 1:200), p-Smad3 Ser423/425 (Abcam ab52903; IHC 1:200), Fibronectin (BD 610078; IHC 1:400), α-Smooth Muscle Actin (Sigma A2547; IHC 1:400). CDK2 (Cell Signaling 2546S; IB 1:1000).

### Statistical analysis

Statistical analysis was performed using GraphPad Prism software (GraphPad software Inc.). Data were analyzed for normality and equal variance using the Shapiro-Wilk test and Brown-Forsythe test, respectively. If both tests were passed, statistical significance was further analyzed by unpaired t test for two groups comparison or one-way ANOVA with Tukey’s post hoc test for multiple groups comparison. A P value less than 0.05 was considered significant (*P < 0.05, **P < 0.01, ***P < 0.001).

## Author contributions

H.D. performed most of the experiments and analyzed data. O.S.R., P.J. and A.T. performed RNAseq and cSTAR analysis. H.D. and O.S.R. prepared figures. H.D. and D.J. prepared all samples for RNAseq. H.X. helped with mouse pressure measurement. B.N.K. and M.A.S. supervised and supported the project. H.D., O.S.R., B.N.K. and M.A.S. wrote the manuscript.

## Acknowledgments

This work was supported by NIH Grants R01 HL135582 (MAS), NIH/NCI grant R01CA244660 and EU grant 101136926 MULTIR (BNK), and SFI grant 22/PATH-S/10875 (OSR).

