## Supplemental Figure 1-2 and Table 1 for "cSTAR analysis identifies endothelial cell cycle as a key regulator of flow-dependent artery remodeling"

**This PDF file incudes:**

**Supplemental Figure Legends**

**Figures S1-3**

**Table S1**

#### Suppl Figure Legends

**Figure S1. The DPD scores for EC drug responses.** (A, B) Dose responses of the $DPD_{remod}$ (A) and the $DPD_{OSS}$ (B) scores to inhibitors of MELK (OTS-167), CDK1/2 (CGP-60474), PAK (PF-03758309), BET bromodomain (JQ1), MEK (PD-0325901), ITK (BMS-509744), p38 (TAK-715), PHD2 (IOX2), and RAF (RAF-265).

**Figure S2. Lung histology images**. (A) H&E staining of Ctrl and CDK2 iECKO lung sections. V, vessels. Arrowheads are narrowed and occlusive vessels.

**Figure S3. Activation of Smad2/3 in CDK2 ECKO lung tissue**. (A-B) Representative immunostaining and quantification of p-Smad2 Ser465/467 (A) and p-Smad3 Ser423/425 (B) in lung sections from Ctrl and CDK2 iECKO mice. Scale bar: 25µm, n=6 mice per group. ***$P$ < 0.001, calculated by two-tailed unpaired t tests.

**Table S1. cSTAR analysis of HUVEC cell state transitions.** Sheet 1. Local response matrix: mean values of the BMRA reconstructed local response matrix elements; Sheet 2. Local response matrix confidence intervals: confidence intervals of mean values of the BMRA reconstructed local response matrix elements; Sheet 3. Global response matrix: cSTAR predicted global response matrix.

### Figure S1

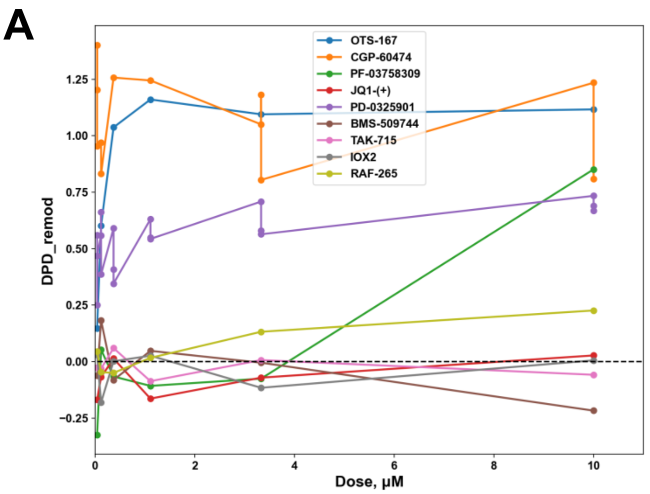
A
DPD_remod
Dose, μM
OTS-167
CGP-60474
PF-03758309
JQ1-(+)
PD-0325901
BMS-509744
TAK-715
IOX2
RAF-265

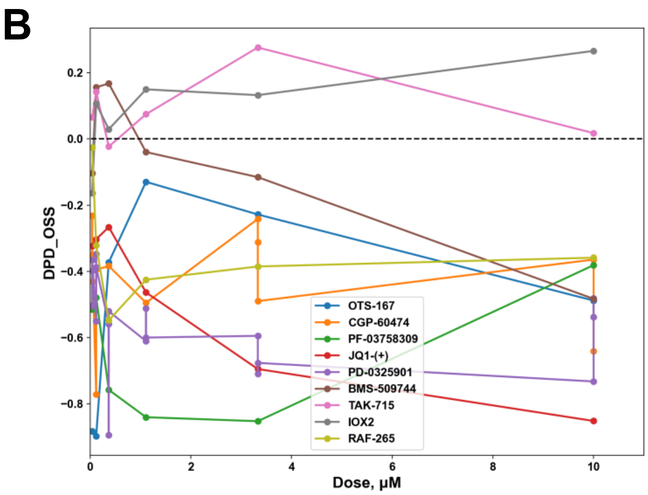
B
DPD_OSS
Dose, μM
OTS-167
CGP-60474
PF-03758309
JQ1-(+)
PD-0325901
BMS-509744
TAK-715
IOX2
RAF-265

### Figure S2

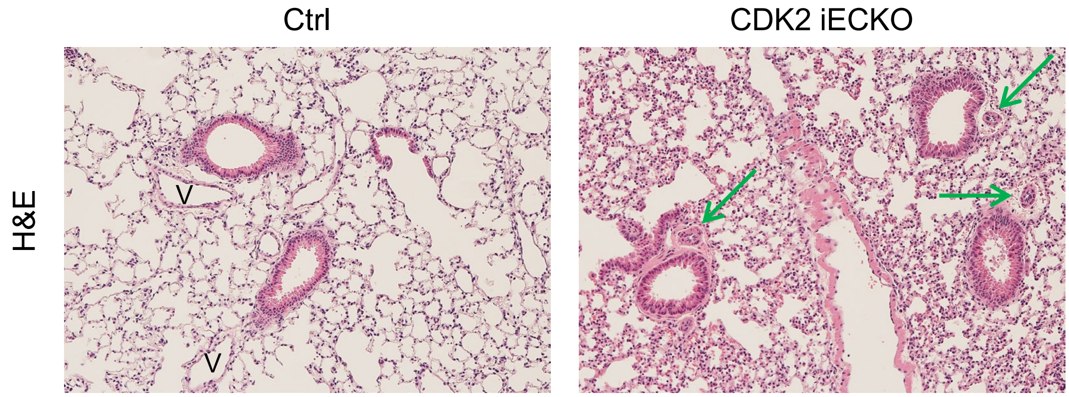

### Figure S3

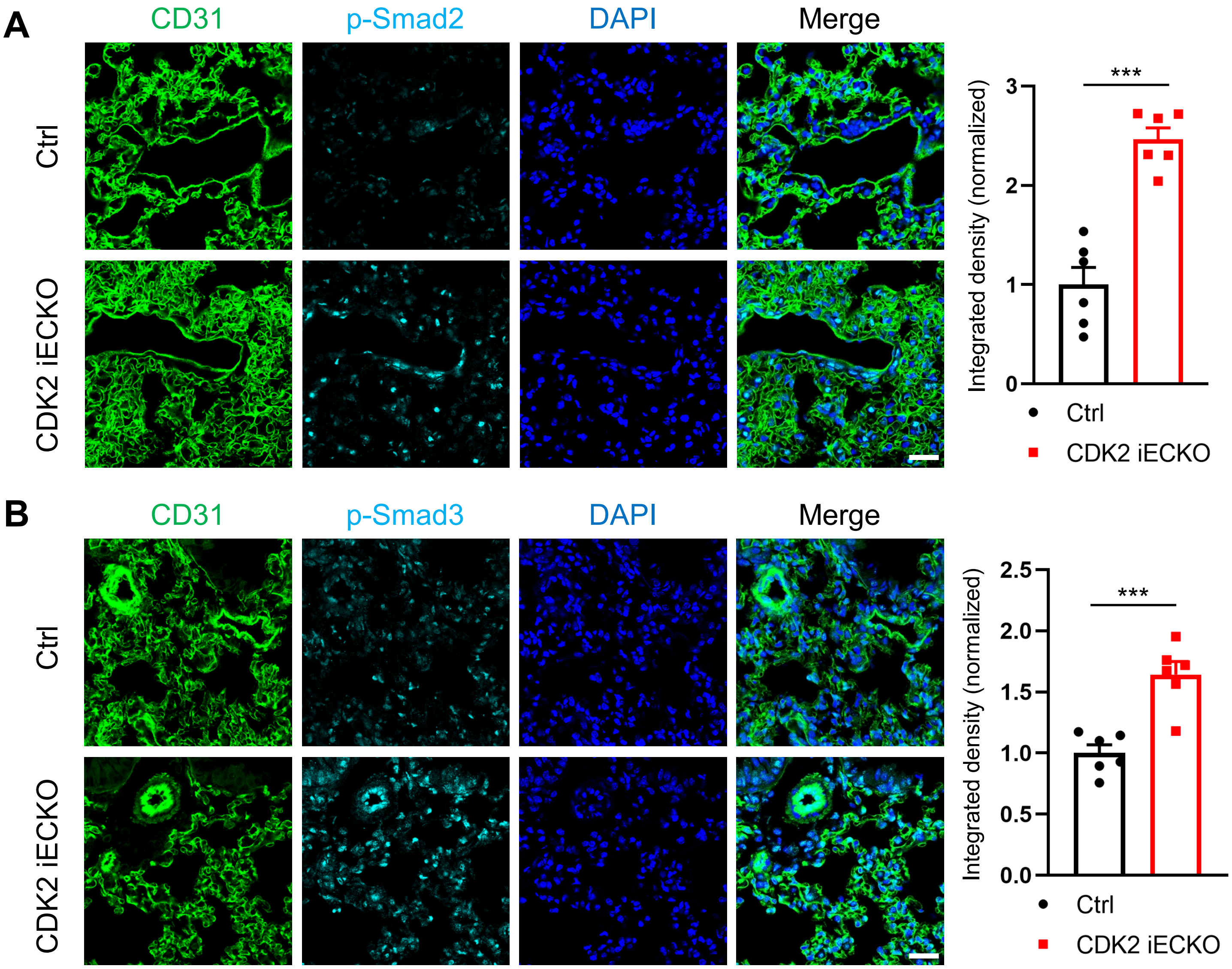
A
CD31
p-Smad2
DAPI
Merge
Ctrl
CDK2 iECKO
Integrated density (normalized)
***
0
1
2
3
Ctrl
CDK2 iECKO
B
CD31
p-Smad3
DAPI
Merge
Ctrl
CDK2 iECKO
Integrated density (normalized)
***
0.0
0.5
1.0
1.5
2.0
2.5
Ctrl
CDK2 iECKO

| **Table S1. cSTAR analysis of HUVEC cell state transitions.** | | |
|---|---|---|
| **Contents** | | |
| | **Sheet name** | **Description** |
| | local response matrix | Mean values of the BMRA reconstructed local response matrix elements |
| | local response matrix confidence intervals | Confidence intervals of the mean values of the BMRA reconstructed local response matrix |
| | Global response matrix | cSTAR predicted global response matrix |

| | CDK1/2 | PAK | PI3K/AKT | MEK/ERK | ITK | p38 | PHD2 | DPD_FSS | DPD_remod | DPD_OSS |
|---|---|---|---|---|---|---|---|---|---|---|
| **CDK1/2** | -1 | 0.184466159 | 0.000583006 | 0.113791472 | 0 | 0 | 0 | 0 | 0 | 0 |
| **PAK** | 0.368922267 | -1 | 0.195858552 | -0.227886072 | 0 | 0 | 0 | 0 | 0 | 0 |
| **PI3K/AKT** | 0.362604792 | 0 | -1 | 0.64306567 | 0.129126909 | 0 | 0 | 0 | 0 | 0 |
| **MEK/ERK** | 0.452922457 | 0 | 0 | -1 | 0 | 0 | 0 | 0 | 0 | 0 |
| **ITK** | -0.370465822 | 0 | 0.358852628 | 0 | -1 | 0 | 0 | 0 | 0 | 0 |
| **p38** | 0.252781 | 0 | 0 | -0.299120635 | 0 | -1 | 0.070246305 | 0 | 0 | 0 |
| **PHD2** | 0 | 0 | 0 | 0.224027637 | 0 | 0.189435846 | -1 | 0 | 0 | 0 |
| **DPD_FSS** | -0.505601945 | 0.268281158 | 0.113474435 | 0.014207925 | 0.054845833 | -0.081353692 | 0 | -1 | 0 | 0 |
| **DPD_remod** | -0.46453949 | 0.10621412 | -0.106504134 | -0.173334571 | 7.34686E-05 | -0.031902625 | 0 | 0 | -1 | 0 |
| **DPD_OSS** | 0 | 0.275418855 | 0.088493524 | 0.198736381 | 0 | -0.091071425 | 0 | 0 | 0 | -1 |
| **Local network impact** | 0.293780424 | 0.016871557 | 0.080340743 | 0.272035987 | 0.008302414 | 0.01778482 | 0.002464235 | | | |
| **Local phenotypic impact** | 0.46453949 | 0.295189744 | 0.138471059 | 0.263706319 | 7.34686E-05 | 0.096497574 | 0 | | | |

| | CDK1/2 | PAK | PI3K/AKT | MEK/ERK | ITK | p38 | PHD2 | DPD_FSS | DPD_remod | DPD_OSS |
|---|---|---|---|---|---|---|---|---|---|---|
| **CDK1/2** | 0 | 0.004243758 | 0.003987037 | 0.003878287 | 0 | 0 | 0 | 0 | 0 | 0 |
| **PAK** | 0.011485875 | 0 | 0.009731362 | 0.011743972 | 0 | 0 | 0 | 0 | 0 | 0 |
| **PI3K/AKT** | 0.030842667 | 0 | 0 | 0.029272301 | 0.069459644 | 0 | 0 | 0 | 0 | 0 |
| **MEK/ERK** | 0.000754571 | 0 | 0 | 0 | 0 | 0 | 0 | 0 | 0 | 0 |
| **ITK** | 0.053315968 | 0 | 0.041440236 | 0 | 0 | 0 | 0 | 0 | 0 | 0 |
| **p38** | 0.039878418 | 0 | 0 | 0.042768136 | 0 | 0 | 0.113365629 | 0 | 0 | 0 |
| **PHD2** | 0 | 0 | 0 | 0.014726285 | 0 | 0.01782737 | 0 | 0 | 0 | 0 |
| **DPD_FSS** | 0.016977799 | 0.01800268 | 0.014167824 | 0.024673919 | 0.009407679 | 0.012747673 | 0 | 0 | 0 | 0 |
| **DPD_remod** | 0.017488157 | 0.01759426 | 0.011750757 | 0.016991066 | 0.001364936 | 0.031898614 | 0 | 0 | 0 | 0 |
| **DPD_OSS** | 0 | 0.012753604 | 0.009810395 | 0.011372468 | 0 | 0.01054531 | 0 | 0 | 0 | 0 |

Global response matrix

| | CDK1/2 | PAK | PI3K/AKT | MEK/ERK | ITK | p38 | PHD2 | DPD_FSS | DPD_remod | DPD_OSS |
|---|---|---|---|---|---|---|---|---|---|---|
| **CDK1/2** | 1.141399308 | 0.210549546 | 0.043939416 | 0.110156128 | 0.005673761 | 1.43362E-16 | -1.70373E-17 | 8.80643E-17 | 2.16972E-16 | -1.1736E-16 |
| **PAK** | 0.445340147 | 1.082150186 | 0.222518985 | -0.052836725 | 0.028733189 | 3.36506E-17 | 2.1485E-16 | 3.8419E-16 | 7.15305E-16 | -1.81068E-16 |
| **PI3K/AKT** | 0.725328225 | 0.133798512 | 1.07651133 | 0.744312828 | 0.13900658 | -1.368E-16 | 3.66883E-16 | 1.13664E-16 | 2.50014E-16 | 9.7375E-17 |
| **MEK/ERK** | 0.51696538 | 0.095362618 | 0.019901148 | 1.049892184 | 0.002569774 | -2.80087E-16 | 1.06688E-16 | 1.52018E-16 | -1.28988E-16 | -1.00662E-17 |
| **ITK** | -0.162563493 | -0.029987463 | 0.370030868 | 0.226289534 | 1.047780942 | 4.87636E-16 | 1.24903E-16 | -6.89736E-16 | 4.95808E-16 | -5.09206E-16 |
| **p38** | 0.143940023 | 0.026552063 | 0.005541129 | -0.273313801 | 0.000715509 | 1.013486637 | 0.071193691 | 9.56664E-17 | -1.44788E-17 | -5.40471E-17 |
| **PHD2** | 0.143081932 | 0.026393774 | 0.005508096 | 0.183429434 | 0.000711243 | 0.191990699 | 1.013486637 | -1.60686E-16 | 1.30844E-17 | -4.16975E-16 |
| **DPD_FSS** | -0.388592106 | 0.196599065 | 0.179764926 | 0.064153142 | 0.078058322 | -0.08245088 | -0.00579187 | 1 | -1.00816E-16 | 9.00513E-17 |
| **DPD_remod** | -0.654386075 | -0.014497965 | -0.11502899 | -0.309302829 | -0.014779869 | -0.032332884 | -0.002271266 | -1.52534E-15 | 1 | 6.09669E-16 |
| **DPD_OSS** | 0.27647293 | 0.326418755 | 0.160000649 | 0.284857485 | 0.020660389 | -0.092299672 | -0.006483711 | -9.0698E-16 | 4.5384E-16 | 1 |
| **Global network impact** | 0.536910058 | 0.115650976 | 0.160905662 | 0.353053961 | 0.057370863 | 0.031511315 | 0.015984107 | | | |
| **Global phenotypic impact** | 0.710393142 | 0.326740561 | 0.197058053 | 0.420490222 | 0.025402681 | 0.097799002 | 0.006870019 | | | |
